# *Oa*AEP1-mediated PNA-protein conjugation enables erasable imaging of membrane protein

**DOI:** 10.1101/2021.11.07.467647

**Authors:** Zhangwei Lu, Yutong Liu, Yibing Deng, Bin Jia, Xuan Ding, Peng Zheng, Zhe Li

## Abstract

Methods to efficiently and site-specifically conjugate proteins to nucleic acids could enable exciting application in bioanalytics and biotechnology. Here, we report the use of the strict protein ligase to covalently ligate a protein to a peptide nucleic acid (PNA). The rapid ligation requires only a short N-terminal GL dipeptide in target protein and a C-terminal NGL tripeptide in PNA. We demonstrate the versatility of this approach by conjugating three different types of proteins with a PNA strand. The biostable PNA strand then serves as a generic landing platform for nucleic acid hybridization. Lastly, we show the erasable imaging of EGFR on HEK293 cell membrane through toehold-mediated strand displacement. This work provides a controlled tool for precise conjugation of proteins with nucleic acids through an extremely small peptide linker and facilitates further study of membrane proteins.

**TOC:** 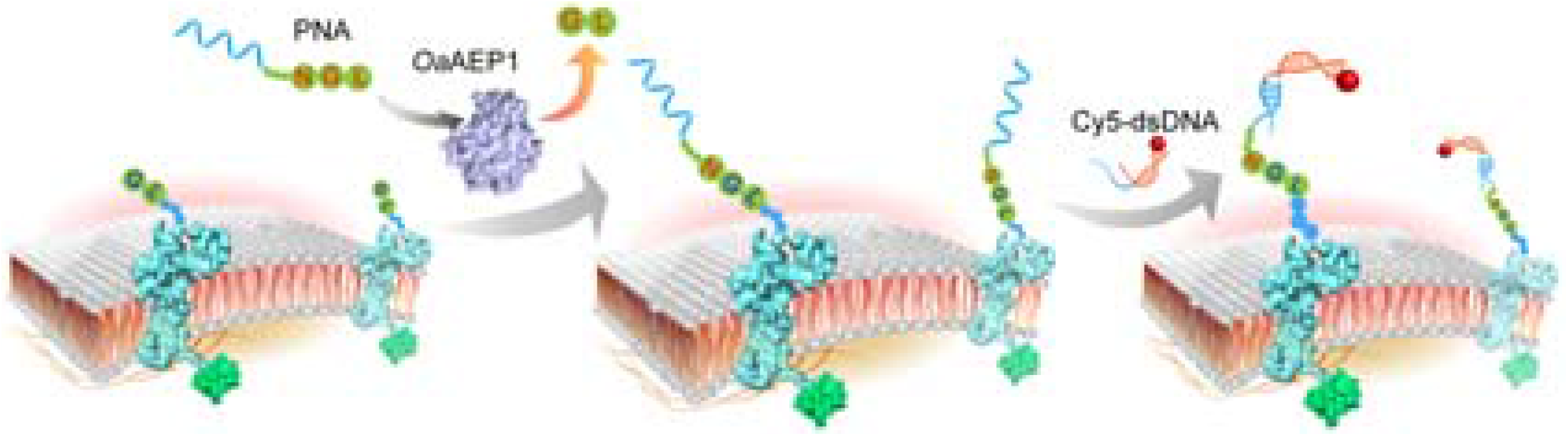

## Introduction

In recent years, the emerging DNA nanotechnology has demonstrated powerful capability to construct addressable and programmable nanostructures, which has served as a reliable platform to manipulate and study proteins at nanometer scale for many biological and biomedical applications^1,2^. For example, DNA origami has been used to precisely control the interenzyme spacing and position of glucose oxidase/horseradish peroxidase enzyme pairs to enhance its activity ^3^. Besides, the spatial presentation of a clinical vaccine immunogen eOD-GT8 has been systematically studied on DNA origami to enhance B-cell activation, offering important foundations for the rational design of molecular vaccines ^4^.

Key to successful application of DNA nanotechnology in protein manipulation and study is a robust method to anchor a nucleic acid strand to a protein of interest (POI), either noncovalently or covalently. A frequently used approach to achieve this relies on DNA aptamers^5,6^, which binds to POI with high affinity and specificity via multiple noncovalent interactions. This approach does not require chemical modifications on either POI or DNA but is only applicable to a small portion of proteins with high-quality aptamer sequences available^7^. Another method involves protein engineering. The POI can be site-specifically modified with an azido group^8^ or a specific peptide tag^9^. Despite high specificity and stability, the method either requires complicated bioorthogonal chemistry or induces a large payload^10^. Therefore, an efficient and precise method to create nucleic acid-protein conjugates is urgently needed.

Recently, an engineered strict protein ligase *Oa*AEP1 (C247A) (*Oldenlandia affinis* asparaginyl endopeptidases 1 with Cys247Ala mutation, abbreviated as *Oa*AEP1 in this paper) was reported^11,12^. As an efficient and strict endopeptidase, *Oa*AEP1 can rapidly link two peptides or proteins covalently by forming a peptide bond through the two termini. Moreover, *Oa*AEP1 requires only a short N-terminal GL dipeptide (NH_2_-Gly-Leu) on one substrate and a C-terminal NGL sequence (Asn-Gly-Leu-COOH) on the other^12,13^. We thus envisioned that a nucleic acid-protein conjugate could be prepared by *Oa*AEP1 from substrates with appropriate peptide modifications. Development of such conjugation methodology would open promising opportunities for the manipulation of POIs with DNA nanotechnology and offer exciting applications in many areas.

Here, we introduce a method that precisely conjugates a POI with a PNA by *Oa*AEP1-mediated enzymatic ligation (Figure 1). Specifically, the POI contains a GL dipeptide at the N-terminus, and the PNA carries a C-terminal NGL tripeptide modification. *Oa*AEP1-catalyzed reaction results in a POI-PNA conjugate with a 3-amino acid linker (NGL). The biostability and programmability of PNA make it a versatile handle to manipulate and study POI. In this work, PNA is first anchored to epidermal growth factor receptor (EGFR) through *Oa*AEP1-catalyzed ligation and subsequent hybridization with a fluorescent DNA probe enables direct visualization of EGFR on living cell surface within 15 minutes. Furthermore, toehold-mediated strand displacement allows the removal of the probe and thus an erasable fluorescent imaging of the target protein.

**Figure 1.**
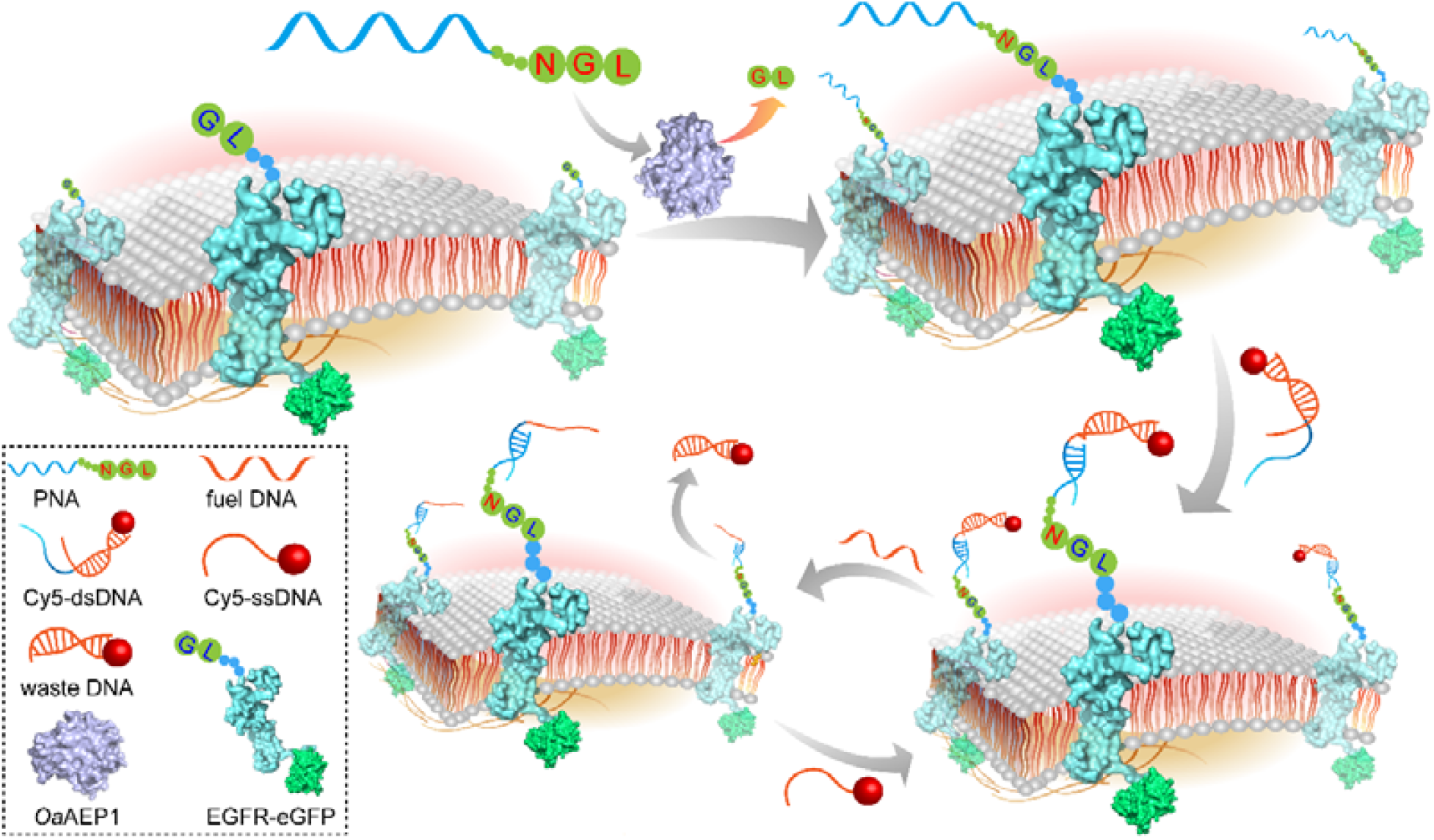
Schematic diagram of erasable imaging of cell membrane proteins. *Oa*AEP1 catalyzes the ligation between EGFR and PNA, which then hybridizes with Cy5-labeled DNA to enable fluorescent imaging. Toehold-mediated strand displacement by fuel strand reverses the labeling process.

## Results and discussion

### PNA-Protein Ligation and Characterization in Solution

To demonstrate the feasibility of *Oa*AEP1-mediated ligation between nucleic acids and proteins, we first modified the 3’ end of a DNA strand with an NGL tripeptide and engineered the N-terminus of protein G containing an additional GL dipeptide. However, *Oa*AEP1-catalyzed ligation only yielded a small amount of product (Figure S1), probably because the negatively charged DNA was not a suitable substrate for *Oa*AEP1. We then considered the electroneutral peptide nucleic acid (PNA). With unique properties, PNA has been used in diverse biological applications, such as genotyping single nucleotide polymorphism (SNP) of human DNA apolipoprotein E gene^14^ and monitoring the internalization of epidermal growth factor receptor (EGFR)^9^. We reason that, given its pseudo-peptide backbone feature and programmable base pairing property, PNA could potentially serve as a substrate for *Oa*AEP1 (Figure 2A) and an adapter molecule between protein and DNA. Finally, the hybridization efficiency of PNA/DNA is stronger than that of DNA/DNA^15^ (Figure S2). Therefore, short PNA strands (10-15 nt) are generally enough to form a specific and stable PNA/DNA hybrid duplex.

**Figure 2.**
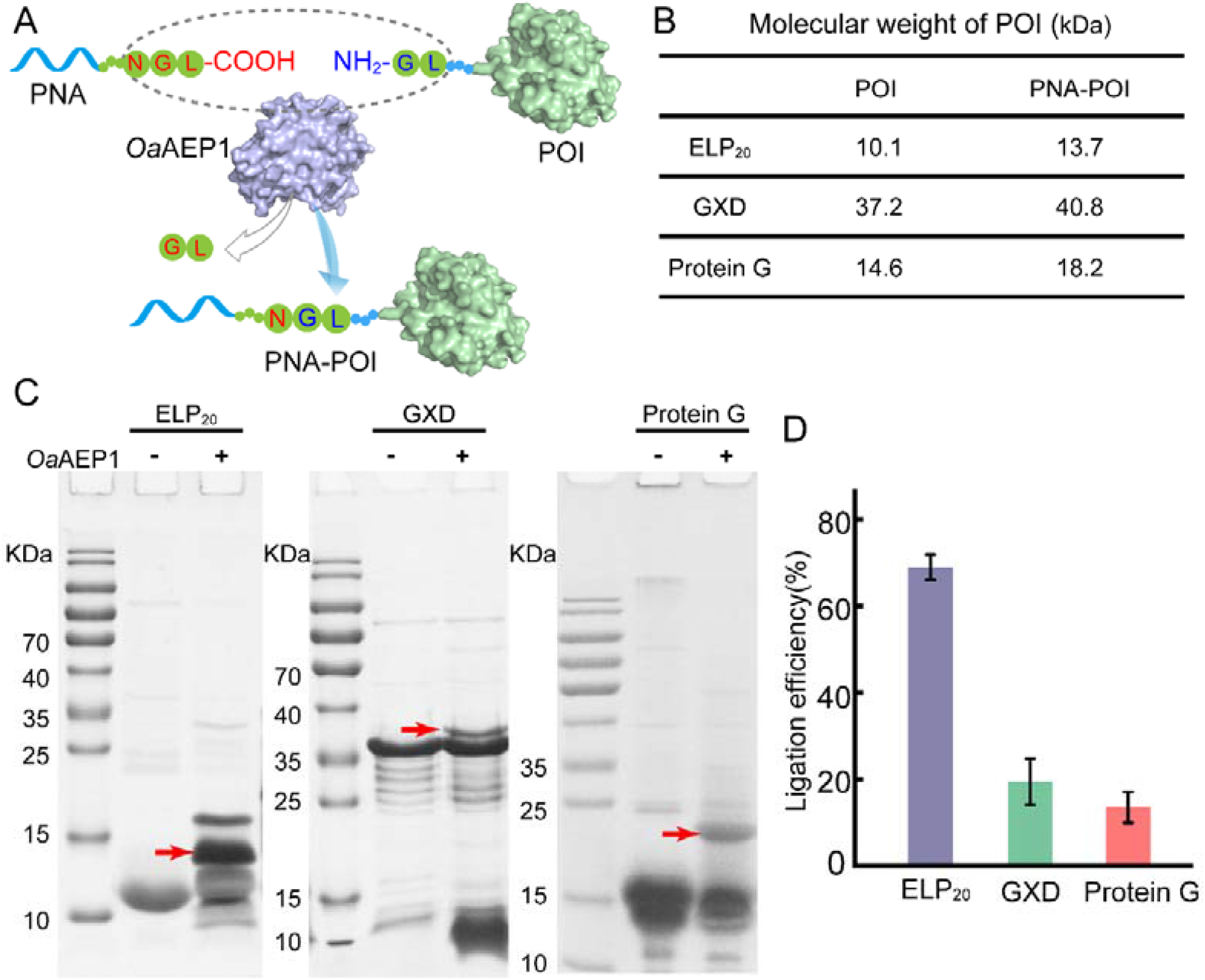
Characterization of PNA-protein ligation catalyzed by *Oa*AEP1. (A) Schematic illustration of the *Oa*AEP1-mediated ligation between PNA and POI. *Oa*AEP1 is in its C247A mutation form (PDB: 4AWA). Protein G (PDB: 1FCC) represents the POI. (B) Molecular weights of three POIs and POI-PNA conjugates. (C) SDS-PAGE analysis of PNA-POI ligation. Red arrows indicate ligation products. (D) Quantitative analysis of the ligation efficiency of each POI with PNA. At least three independent experiments were performed for each protein.

A 12-mer PNA strand modified with a C-terminal NGL tripeptide and a two amino acid linker (Gly-Gly) in between was synthesized (PNA-2aa, 3.61 kDa, Table S1). Simultaneously, three different POIs each containing a GL dipeptide at the N-terminus were expressed (Table S2), including elastin-like polypeptides (ELP_20_)^16^, protein G^17^, and the engineered multidomain protein GB1-X module-Dockerin (abbreviated as GXD)^18^. The molecular weights of these proteins ranged from 10 to 40 kDa (Figure 2B). SDS-PAGE analysis of *Oa*AEP1-catalyzed PNA-POI ligation clearly showed a mobility shift, consistent with the calculated molecular weights of conjugates (Figure 2C). Together with the MALDI mass spectrometry verification (Figure S3), the results not only validated the covalent linkage between PNA and POI, but also demonstrated the general applicability of the method. The ligation efficiency of this one-step reaction between each POI and PNA was then examined (Figure 2D). Interestingly, ELP_20_, which has a flexible structure, showed the highest efficiency of up to ∼70%, while the other two proteins, which have compact globular structures, showed a lower efficiency of less than 20%.

### PNA-Protein Ligation and Characterization on DNA Origami Surface

Given the feasibility of conjugating PNA to POI with high efficiency, as well as the programmable PNA/DNA interaction^19^ we envisioned that this reaction could be conducted on the surface of DNA origami nanostructures and thus enable the site-specific organization of proteins. To achieve this goal, two approaches could be executed. In the ligation-first approach, POI and PNA were first ligated and then the conjugate was installed onto DNA origami nanostructure by PNA/DNA hybridization (Figure 3A). Whereas in the hybridization-first approach, the PNA strand was first immobilized on the DNA origami nanostructure, and then *Oa*AEP1 catalyzed the PNA-POI ligation, positioning POI precisely onto the origami (Figure 3B). In the following experiments, a classic triangular origami nanostructure with edge length of approximately 120 nm was utilized (Figure S4). Custom-selected staple strands were extended with complementary sequences to PNA to serve as capture handles (Table S4).

**Figure 3.**
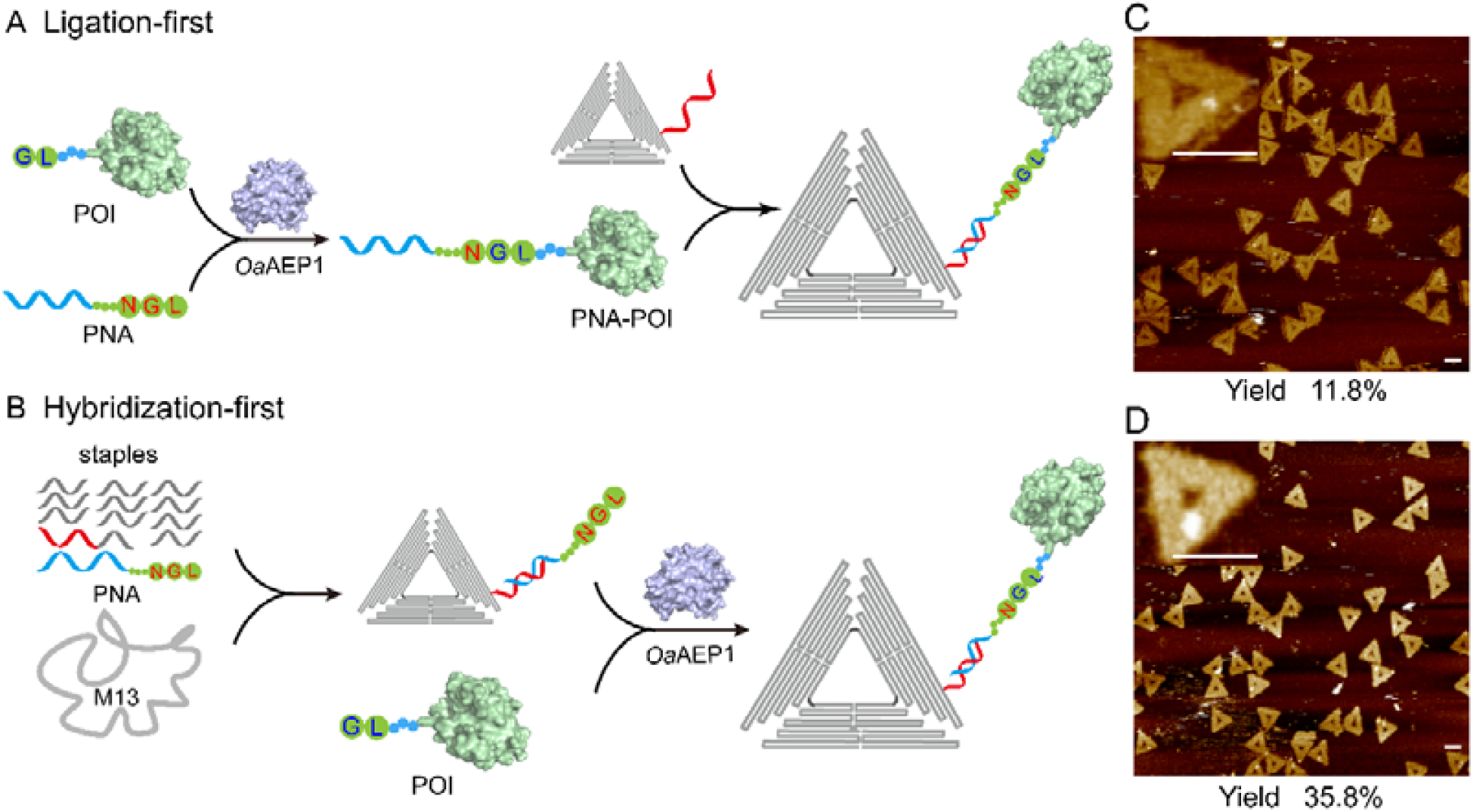
Site-specific organization of POI on DNA origami nanostructures. Schemes of the ligation-first method (A) and hybridization-first method (B). Protein G was organized onto DNA origami triangles using two methods, and the yield was calculated as the percentage of protein G-functionalized nanostructures (C and D). Scale bar: 100 nm.

Protein G could be organized onto the designated sites by both methods. However, the ligation-first method resulted in a lower yield of 11.8% (Figure 3C, calculated as the percentage of protein-functionalized nanostructures), while the hybridization-first method increased the yield to 35.8% (Figure 3D). We reason that in the ligation-first method, the ligation product was difficult to separate from the unreacted substrates. Excess free PNA strand could compete with PNA-POI conjugate for the hybridization opportunities with the capture strands on origami, resulting in a lower yield. Whereas in the hybridization-first method, this problem was alleviated by the removal of excess PNA strands from assembled origami structures using Millipore centrifuge filters. Interestingly, compared with ligation reactions in solution, the ligation yield of PNA and protein G on a DNA origami platform appeared to increase by two fold (35.8% vs. 17.2%). It is worth noting that the yield is calculated as the percentage of protein G-functionalized nanostructures, which takes into account the efficiencies of both POI-PNA ligation and PNA/DNA hybridization, and therefore represents the lower limit of the actual ligation efficiency. We concluded that DNA origami served as a solid support and facilitated the ligation reaction^20^.

We then examined the factors that might affect the reaction yield of POI ligation onto DNA origami nanostructure. We hypothesized that a longer linker within PNA would offer increased flexibility to the terminal NGL tripeptide and thus promote PNA-POI ligation. We thus evaluated the ligation reactions between each of the three POIs and a PNA strand (PNA-6aa, 3.92 kDa) with a six amino acid linker (Gly-Gly-Ser-Gly-Gly-Ser). As expected, compared to the PNA-2aa, PNA-6aa increased the yields for all three POIs (Figure 4A and Figure S5). We inferred that a longer spacer led to less steric hindrance between *Oa*AEP1 enzyme and the PNA substrate. Subsequently, the ligation reaction between POI and PNA-6aa on DNA origami triangles was conducted at pH 5, 6 and 7. AFM images (Figure S6) clearly showed the successful organization of all three POIs on the designated sites with different yields (Figure 4B, Table S3). Generally, acidic solutions (pH 4 to 7) showed little effect on the activity of *Oa*AEP1^12^, however, they might affect the complementary base paring of DNA/DNA^21, 22^ (between staple strands/capture strands and origami scaffold) and of PNA/DNA (Figure S7) (between PNA-ligated POI and capture strands), and therefore, reactions at pH 5 showed less DNA origami triangles in sight and the lowest yield. Finally, we conducted ligation reactions at 10°C and 25°C. The relatively high hybridization efficiency and enzyme activity at room temperature resulted in a higher yield (Figure 4C, Figure S8). It is also worth noting that the yield did not decrease greatly at 10□. Therefore, our method was suitable for the study of temperature-sensitive proteins such as cold shock proteins^23^ (CSPs).

**Figure 4.**
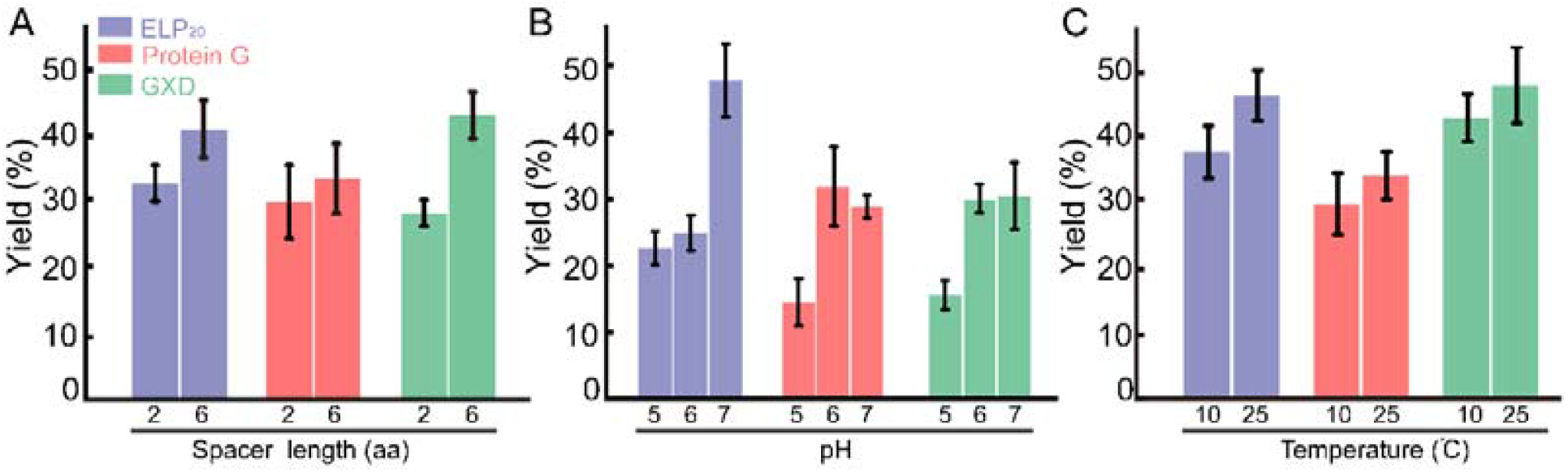
Factors including spacer length (A), pH (B), and temperature (C) affecting *Oa*AEP1-catalyzed ligation yield on DNA origami. Statistical analysis is derived from calculations of 100 tiles.

### Erasable fluorescent imaging of EGFR through *Oa*AEP1-catalyzed ligation, fluorescent DNA probe hybridization and toehold-mediated strand displacement

The genetically encodable terminal GL dipeptide on POI would facilitate the manipulation of membrane proteins on live cells. To demonstrate the feasibility, we carried out the live cell imaging of EGFR (Epidermal Growth Factor Receptor), a transmembrane receptor tyrosine kinase that is often overexpressed in cancer cells and participates in the process of tumor cell proliferation, angiogenesis, tumor invasion, metastasis, and inhibition of apoptosis^24^. EGF binding induces EGFR dimerization and activate signal transduction through autophosphorylation of tyrosine residues, triggering gene transcription and pathways that control cell proliferation and differentiation^25^.

An EGFR plasmid consisting of an N-terminal GL dipeptide and a C-terminal enhanced green fluorescent protein (eGFP) fusion was constructed (Figure S9) and transiently transfected into HEK293 cells (Figure 5A). Confocal laser scanning microscope (CLSM) of eGFP demonstrated that EGFR-eGFP was successfully expressed and correctly localized on the cell membrane (Figure 5B). We then incubated the cells with PNA-6aa and *Oa*AEP1 for 30 minutes to covalently ligate PNA and EGFR. A branched Cy5-dsDNA strand was then added and incubated for 15 minutes (Table S5). The 5’ overhang carries a 12-nt region complementary to PNA, and the 8-nt 3’ overhang serves as a toehold handle for strand displacement. Intense fluorescence signals in both red and green channels were observed on the cell membrane, demonstrating the effective ligation by *Oa*AEP1 on cell surface and the subsequent illumination of target protein with a fluorescent DNA probe. On the contrary, without *Oa*AEP1, the Cy5 signal on the cell surface was negligible (Figure 5C).

**Figure 5.**
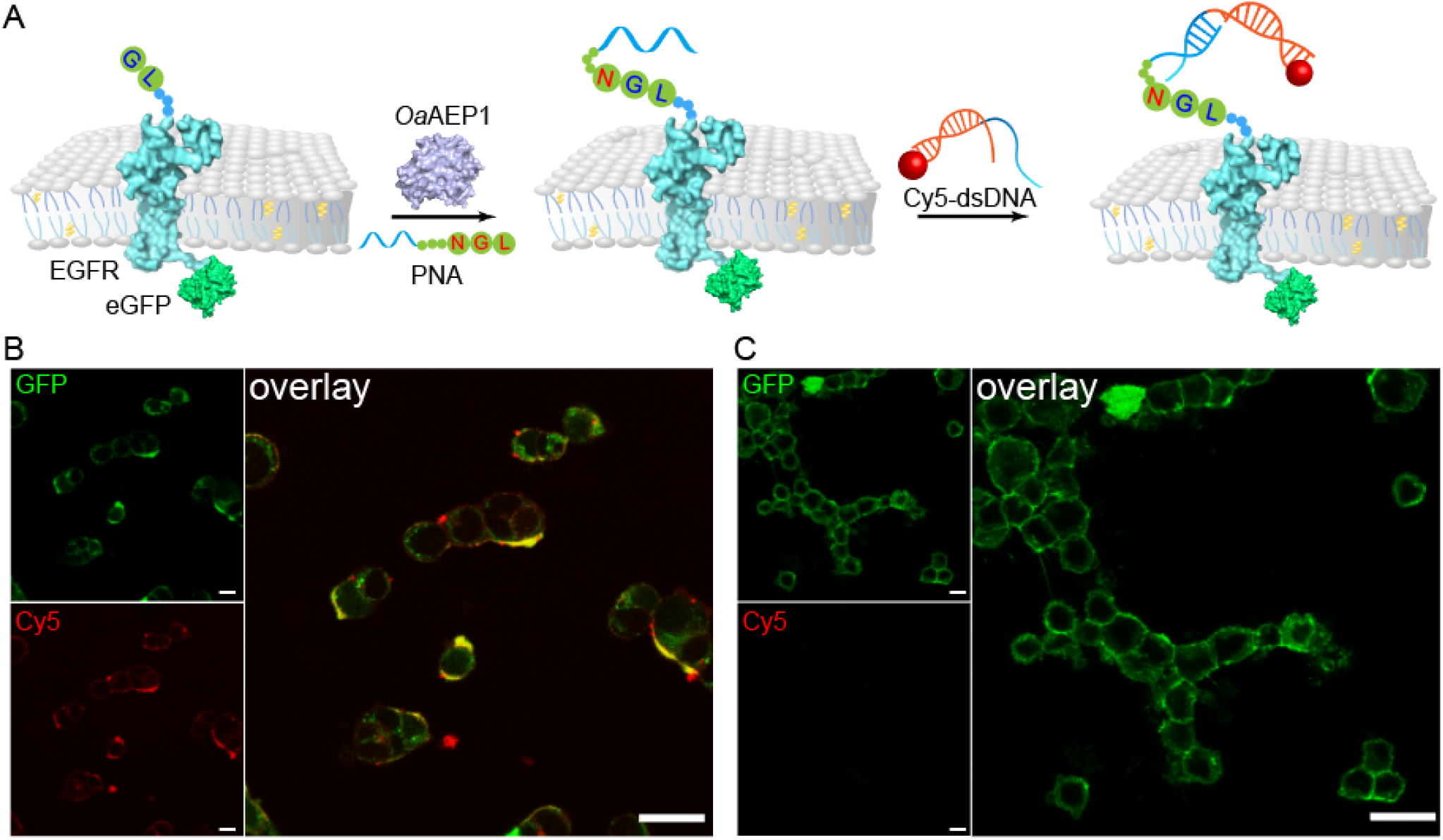
Fluorescent imaging of cell membrane proteins. (A) Schematic diagram of EGFR imaging enabled by PNA ligation and Cy5-DNA hybridization. CYSM images of EGFR on the HEK293 cell surface with (B) or without (C) the incubation with *Oa*AEP1. Fused eGFP was used as an internal control. Scale bars represent 20 μm.

Finally, thanks to the programmability of PNA-DNA interactions, we demonstrated the reversible EGFR imaging using the toehold strand displacement method^26^. After *Oa*AEP1-catalyzed ligation and the DNA-assisted labeling, a fuel DNA strand was added to remove the Cy5-labeled DNA probe (Figure 6A). CLSM confirmed that the Cy5 probe was fully displaceable by the fuel strand, achieving reversible membrane protein imaging. With this targeted protein imaging demonstration, we envisioned that this method could provide a valuable tool to study protein function in a temporally controlled manner and to manipulate membrane protein-protein interaction dynamically.

**Figure 6.**
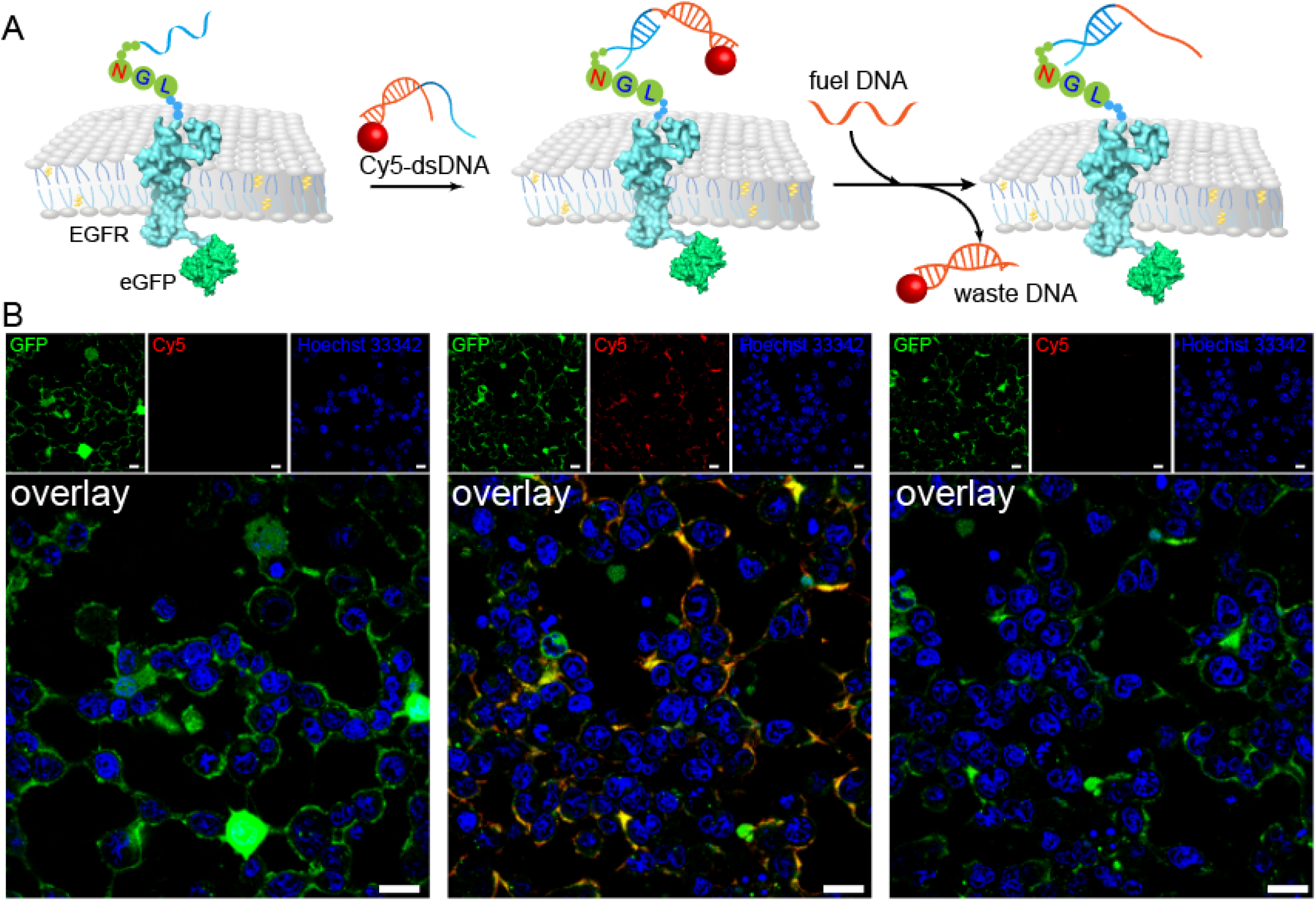
Erasable fluorescent imaging of cell membrane protein EGFR on HEK293 cells. (A) Schematic diagram of erasable fluorescent imaging of EGFR. (B) CYSM images of EGFR on HEK293 cell surface in different conditions: after ligation of PNA (left); after incubation of Cy5-DNA (middle); and after strand displacement (right). Scale bars represent 20 μm.

## Discussion

In this study, we developed a robust enzymatic method for efficient and precise conjugation of proteins to PNA. This ligation is catalyzed by the strict protein ligase *Oa*AEP1 and requires only a short N-terminal GL dipeptide in the target protein and a C-terminal NGL tripeptide in the PNA strand; therefore, it should be applicable for a wide range of proteins. We have demonstrated the versatility of this approach by conjugation of three different types of proteins, ELP_20_, protein G, and GXD, under various conditions. Besides, the resulting linkage between POI and PNA is covalent, which promises stability and therefore is appealing for applications in harsh environment.

In this method, one PNA is fused explicitly to the N-terminus of POI, enabling complete control of the coupling site with minimal possible interference to the protein function. We have also shown that ligation products are observable within minutes and saturated within 2 h. Fast ligation kinetics is especially appealing for many biotechnological applications. In fact, the enzyme *Oa*AEP1 has enabled fast synthesis and immobilization of protein pentamers, and introduced a new generation method of protein preparation/immobilization for AFM-based single-molecule force spectroscopy (AFM-SMFS)^12,27, 28^.

In contrast to DNA, PNA is electroneutral, facilitating its ligation with proteins by *Oa*AEP1. In addition, the hybridization efficiency between PNA/DNA is stronger than that between DNA/DNA^15,29^, and the PNA/DNA duplexes are also resistant to nucleases^30^. The increased thermostability and biostability would further expand the applications of PNA-proteins conjugates. The programmability of PNA also enables manipulation of membrane proteins on the surface of living cells, and thus holds great potential in practical applications. Besides fluorescent imaging of membrane proteins in this study, the use of more complex DNA nanostructures could induce polymerization of receptor proteins on the surface of living cells, and thus provide powerful tools for in-depth study of protein signal transduction and corresponding activation processes.

In conclusion, the enzymatic ligation of proteins with PNA provides a new method to equip proteins with a generically addressable and biostable control unit, and thus may prove useful for a broad range of biotechnological applications.

## Supporting information

Detailed experimental methods, additional gel electrophoresis and AFM images, and oligonucleotide sequences are supplied as Supporting Information.

## Acknowledgment

This work was supported by the Fundamental Research Funds for the Central Universities (Grant No. 14380259), Natural Science Foundation of Jiangsu Province (No. BK20200058), Program for Innovative Talents and Entrepreneur in Jiangsu, Jiangsu Province Key Research and Development Program: Social Development Project (BE2021653), and the National Natural Science Foundation of China (Grant No. 21771103, 21977047). We thank H. Yu for critical reading and commenting on this manuscript.

