## supplemental for "*Oa*AEP1-mediated PNA-protein conjugation enables erasable imaging of membrane protein"

***Oa*AEP1-mediated erasable imaging of membrane proteins through programmable PNA-DNA assembly**

### Materials

All DNA oligonucleotides were purchased from Sangon Biotech Co., Ltd. (Shanghai, China). E. coil BL21 (DE3) and XL1-Blue cells were purchased from TransGen Biotech Co., Ltd (Beijing, China). All other reagents were purchased from Sangon Biotech Co., Ltd. (Shanghai, China) or Sigma-Aldrich (USA). HEK293 cell was purchased from KeyGEN Biotech Co., Ltd.(Nanjing, Jiangsu).

DNA oligonucleotides were dissolved in 1×PBS (1×PB, 300 mM NaCl, pH 7.4) or 1×TAE Mg^2+^ (40 mM Tris, 40 mM Acetic acid, 1 mM EDTA, 12.5 mM Mg^2+^, pH 8.5) and stored at -20 °C for use.

### Protein engineering

ELP is abbreviated for elastin-like polypeptides. X module and Doc are from type III cohesin-dockerin-X module domain complex of Ruminococcus flavefaciens. GB1 is the B1 domain of streptococcal protein G. *Oa*AEP1 (C247A), abbreviated as *Oa*AEP1, is the cysteine 247 to alanine mutation of asparaginyl endoproteases 1 from oldenlandia affinis. All other plasmids were purchased from Genscript Inc. Genes encoding for target proteins, such as GL-ELP20, GL-GXD, and GL-protein G were constructed in expression vector pET-28a by standard molecular biology techniques.

All the plasmids were transformed in *E. coli* BL21 (DE3) for overexpression. The bacteria cultured in LB medium were induced by the addition of 1 mM IPTG (final concentration) overnight at 16 °C. Then, cells were collected by centrifugation at 8000 rpm for 10 min, and resuspended and lysed by high-pressure homogenization. After centrifuged at 18000 rpm for 15 min, the supernatants were loaded onto the Co^2+^ affinity chromatography column. The recombinant proteins were washed with washing buffer (20 mM Tris, 400 mM NaCl, 2 mM imidazole, pH 7.4) and subsequently eluted in elution buffer (20 mM Tris, 400 mM NaCl, 250 mM imidazole, pH 7.4). Finally, the purified proteins were transferred in buffer without imidazole (50 mM Tris, 100 mM NaCl, pH 7.4) through ultrafiltration and can be stored at -80°C after the addition of 20% glycerol (final concentration). The expression and purification of *Oa*AEP1 can be found in literature^1^^,^^2^.

### POI and DNA ligation by *Oa*AEP1

HS-DNA (100 μM, 50 μL) was fist reacted with a 20-fold excess of Sulfo-SMCC in 1× PBS (pH 7) for two hours. Excess SMCC was removed by Amicon, 3KDa cutoff filters. Next, SMCC modified DNA was conjugated to peptide GGNGL. DNA-GGNGL (50 μM, 10 μL) and GL-protein G (20 μM, 10 μL) were mixed with the addition of 1 μM *Oa*AEP1 at 25°C for 2 hours. The reaction was stopped in SDS loading buffer and characterized by SDS-PAGE gel. Only a small amount of ligated products were observed (Figure S1).

### POI and PNA ligation by *Oa*AEP1

PNA-2aa/PNA-6aa (50 μM, 10 μl) and GL-POI (20 μM, 10 μL) were mixed with the addition of 1 μM *Oa*AEP1. Different reaction conditions including spacer length, reaction time, pH and temperature were employed. The reaction was stopped in SDS loading buffer and characterized by SDS-PAGE gel and MALDI mass-spectrometry (Figure S2). The catalytic efficiency for each reaction was calculated based on the band intensity of the reactant/product using the software BandScan.

### Synthesis of DNA origami

A triangular DNA origami (Figure S3) was annealed, referring to Yan's protocol ^4^. Scaffold M13mp18 DNA (5 nM) was mixed with staple strands and PNA-2aa/PNA-6aa in a 1:5 ratio. The DNA origami was assembled in 1×TAE-Mg buffer (40 mM Tris, 40 mM Acetic acid, 1 mM EDTA, 12.5 mM Mg^2+^) by cooling down from 85°C to 4°C. 100 kDa MWCO Amicon filters were used to remove the excess staple and PNA stands.

### PNA-protein conjugation on DNA origami

PNA modified DNA origami triangles (20 nM) and GL-POI (200 nM) were mixed with the addition of 0.5 μM *Oa*AEP1. Different reaction conditions, including spacer length, reaction time and pH were employed. The reaction was stopped in 1×TAE-Mg^2+^ buffer and characterized by atomic force microscope (AFM) (Figure S4-6). The catalytic efficiency for each reaction was calculated based on the band intensity of the reactant/product using the software BandScan.

To acquire AFM image, DNA origami or DNA origami-POI complex (1 nM, 3 μL) diluted with TAE buffer containing 3 mM NiCl_2_ was added onto the surface of the freshly uncovered mica and adsorbed for 3 min. Then, 30 μL TAE buffer was added onto the mica surface. Under the liquid phase model of the AFM, triangle origami or origami-POI complex was imaged.

### Plasmids design and expression of GL-EGFR-eGFP on HEK293 cell surface

A plasmid for GL-EGFR-eGFP was cloned into the egfp-N1 vector by Gibson assembly using a homemade mixture. To locate the fusion protein onto cell membrane, a signal peptide sequence was added in front. The sequence of the GL-KGG peptide was inserted between the signal peptide and EGFR (Figure S7). After expression and cleavage of the signal peptide, the mature protein with N terminal GL-KGG tag should be revealed.

Human embryonic kidney cells (HEK 293) were grown as monolayers at 37 °C, 5 % CO_2_, and 95% humidity if not stated otherwise. Cells were cultured in Dulbecco’s Modified Eagle’s Medium (DMEM) from Thermo Fischer Scientific with additional 2mM L-glutamine, and penicillin / streptomycin (10,000 units/ mL), G418 (50 μg/mL) and 10 % FBS. Plating cells were then washed with PBS and detached with 1 mL 0.25% trypsin / 0.02% EDTA solution for 2 min at 37°C. Cells were centrifuged and cultured in fresh medium.

To generate a stable cell line carrying G418 inducible GL-EGFR-eGFP, HEK293 cells were transfected with the vector at a 3:1 ratio using Lipofectamine 3000 (Thermo Fisher Scientific) according to the manufacturer’s instructions. 6-8 hours after, 5 ml DMEM with 10% serum was added, and the cells were incubated for 72 hours.

### Fluorescence microscopic analysis

The transiently transfected cells were treated with *Oa*AEP1 (1 μM, 50 μL) and PNA (50 nM, 200 μL) for ligation. After washing with PBS, we hybridized the PNA strand with its complementary, fluorescently labelled Cy5-DNA (100 nM, 10 μL for 15 min). After washing with PBS to get rid of excess strands, the cell were stained with Hoechst 33342, and imaged under fluorescence microscope in two different channels (eGFP: λ_ex_ = 490 ± 20 nm, λ_em_ > 520 nm; Cy5: λ_ex_ = 600 ± 50 nm, λ_em_ = 650 ± 30 nm).

Subsequently, toehold-mediated strand displacement of the Cy5-DNA was carried out with 300 nM , 20 μL fuel strand for 10 min in DMEM. Transfected cells were then imaged under fluorescent microscopy in three different channels (Hoechst 33342: λ_ex_ = 350 ± 50 nm, λ_em_ = 460 ± 50 nm; eGFP: λ_ex_ = 490 ± 20 nm, λ_em_ > 520 nm; Cy5: λ_ex_ = 600 ± 50 nm, λ_em_ = 650 ± 30 nm).

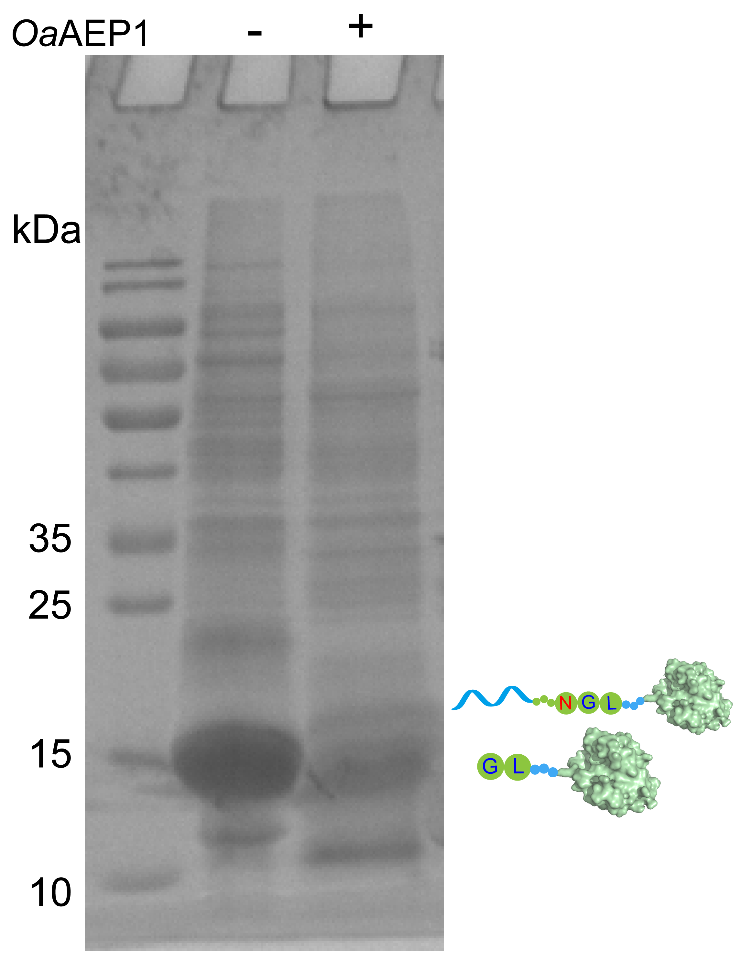

**Figure S1**. SDS-PAGE analysis of protein G (MW 14.6 kDa) before and after ligation with DNA by *Oa*AEP1. The expected molecular weight of the ligation product is 18.9 kDa.

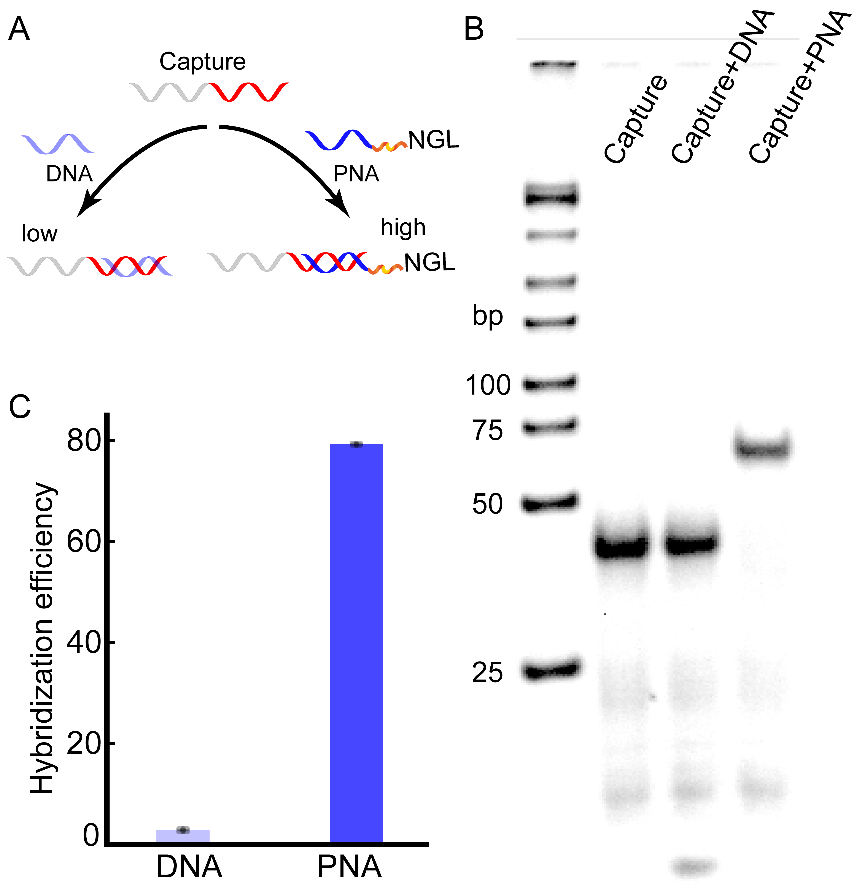

**Figure S2.** Comparison of hybridization efficiency between PNA/DNA and DNA/DNA. (A) Schematic illustration of the hybridization process. DNA capture strand (1 μM) was mixed with PNA-2aa, or a DNA strand with the same sequence, in 1:1 ratio. (B) Native 12% PAGE electrophoresis characterized the hybridization. (C) Gel quantitative analysis of the hybridization efficiency based on the band intensity of the reactant/product using the software BandScan. PNA showed a much higher hybridization efficiency to the capture strand than its DNA counterpart.

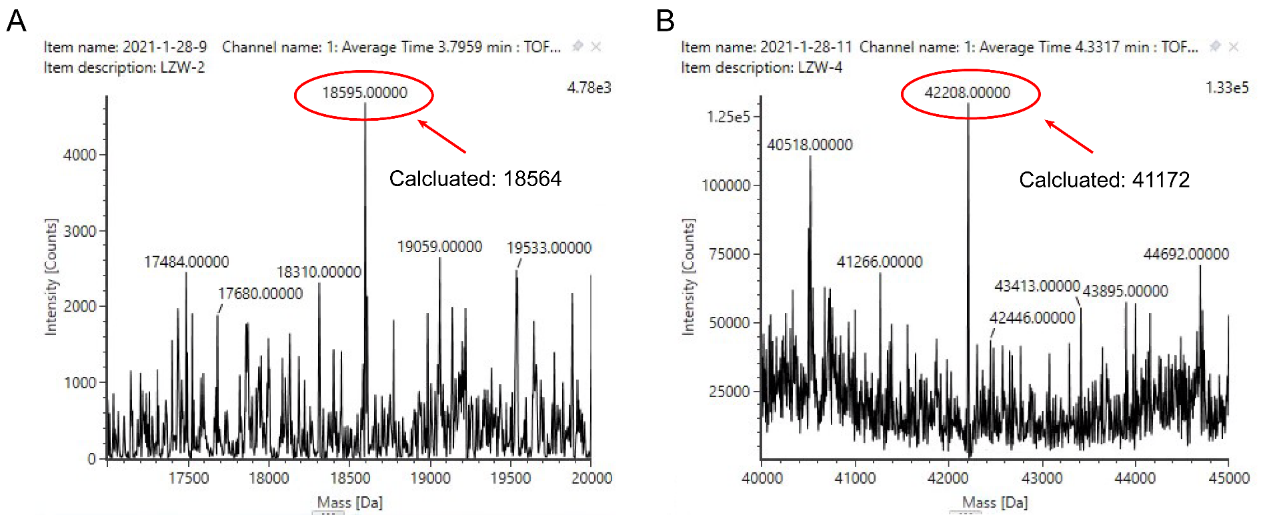

**Figure S3**. MALDI-MS analysis of PNA-protein G (A) and PNA-GXD (B).

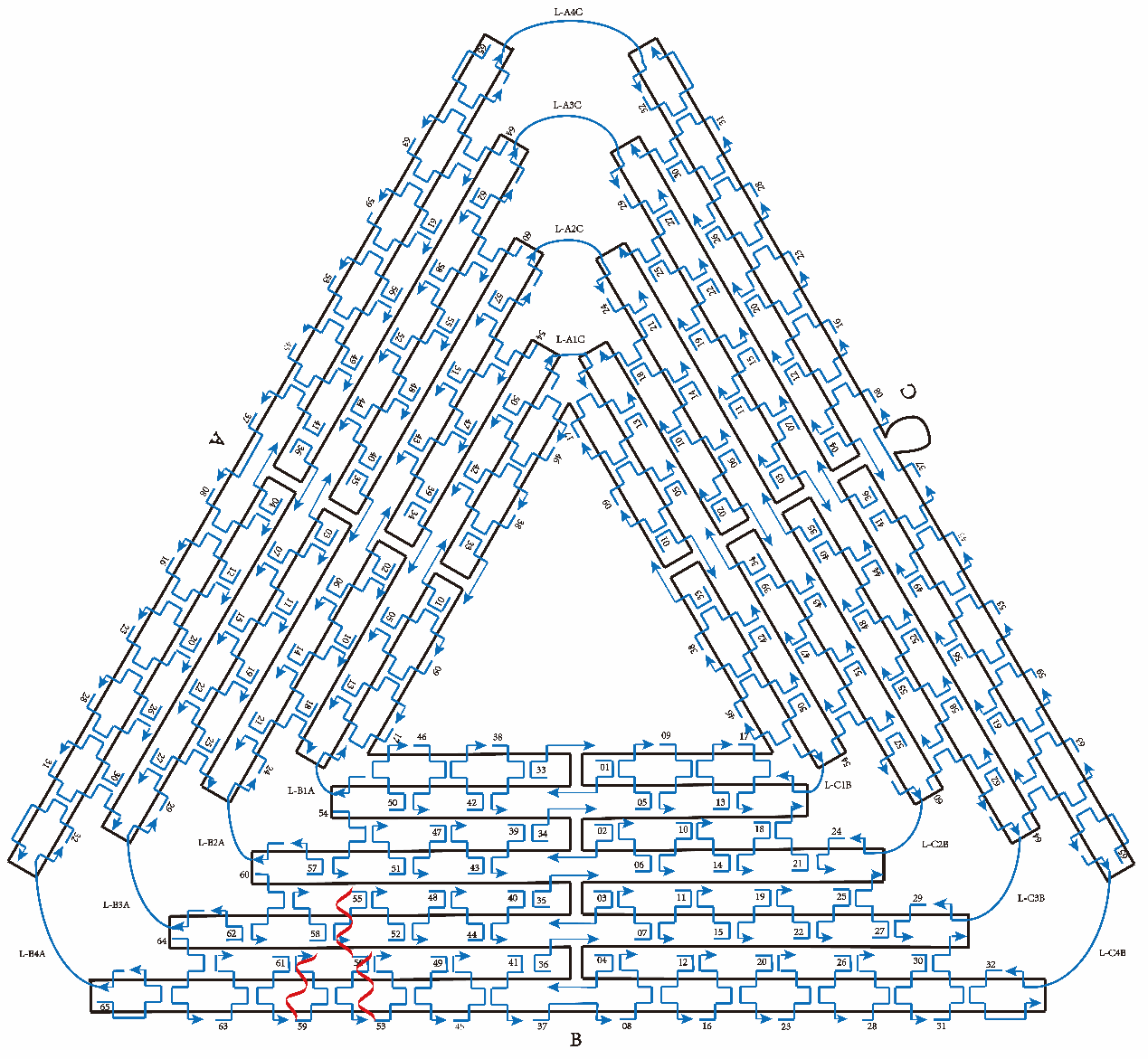

**Figure S4**. Design of the triangular DNA origami. Highlighted in red are capture strands which hybridize with PNA.

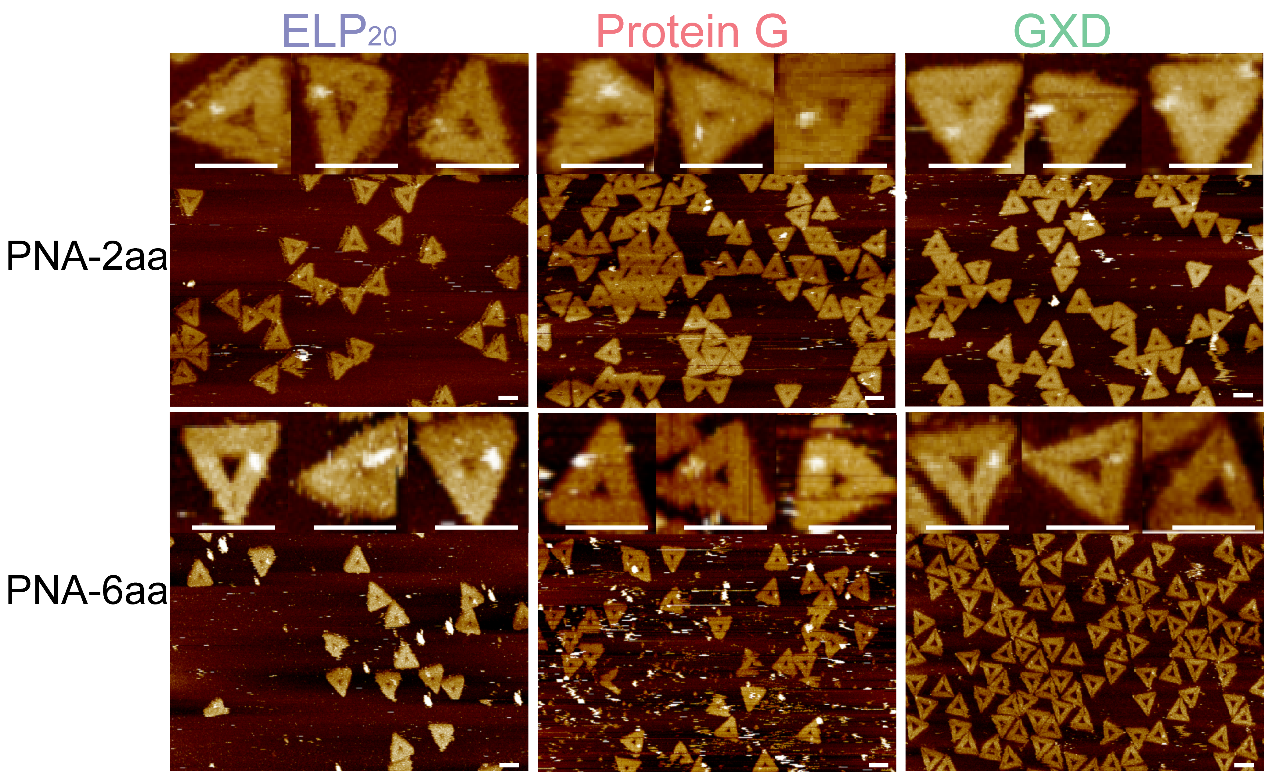

**Figure S5**. AFM images of site-specific organization of ELP_20_, protein G, and GXD through *Oa*AEP1 catalyzed ligation under different spacer lengths. All scale bars: 100 nm.

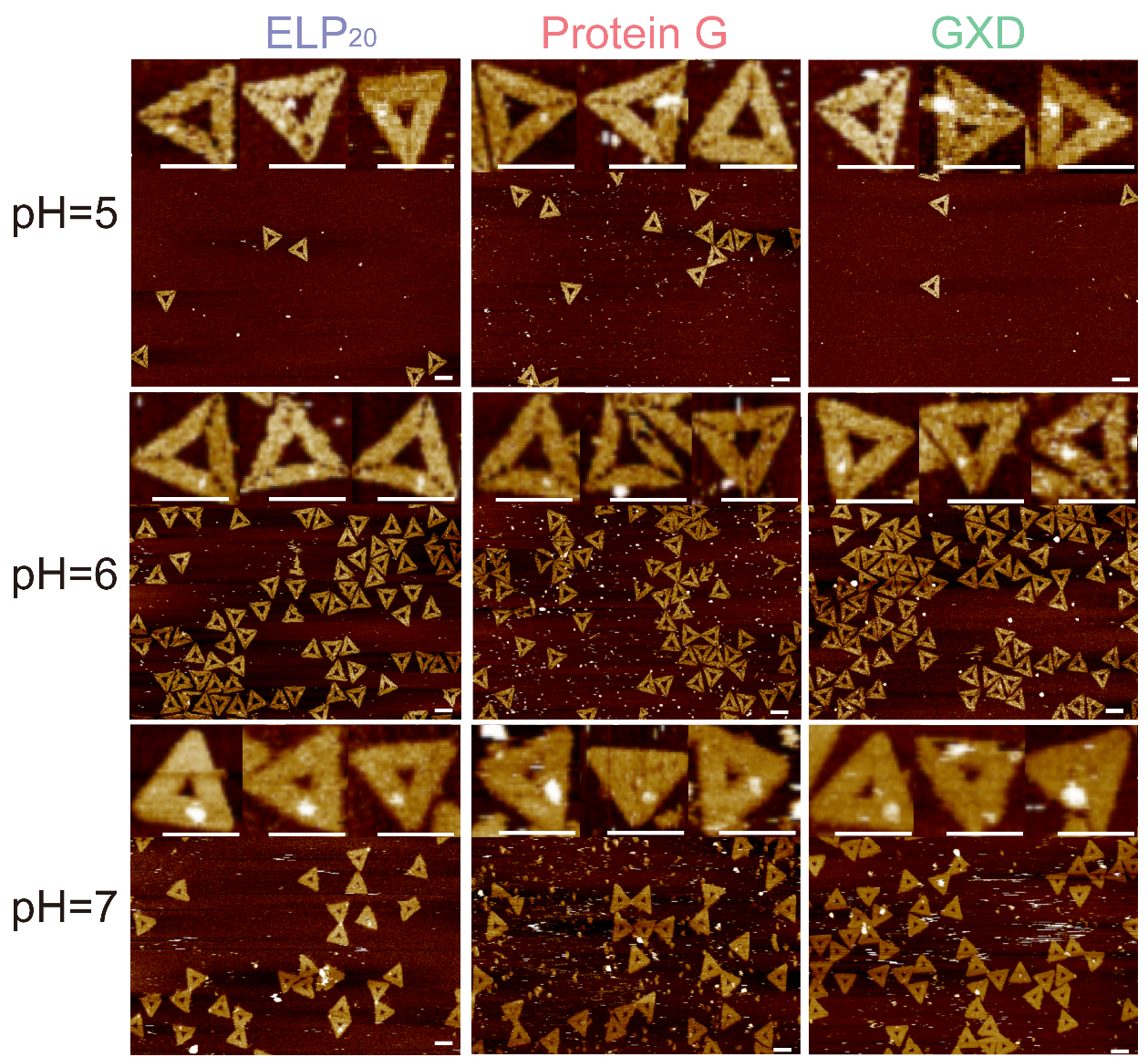

**Figure S6**. AFM images of site-specific organization of ELP_20_, protein G, and GXD through *Oa*AEP1 catalyzed ligation under different pH. All scale bars: 100 nm.

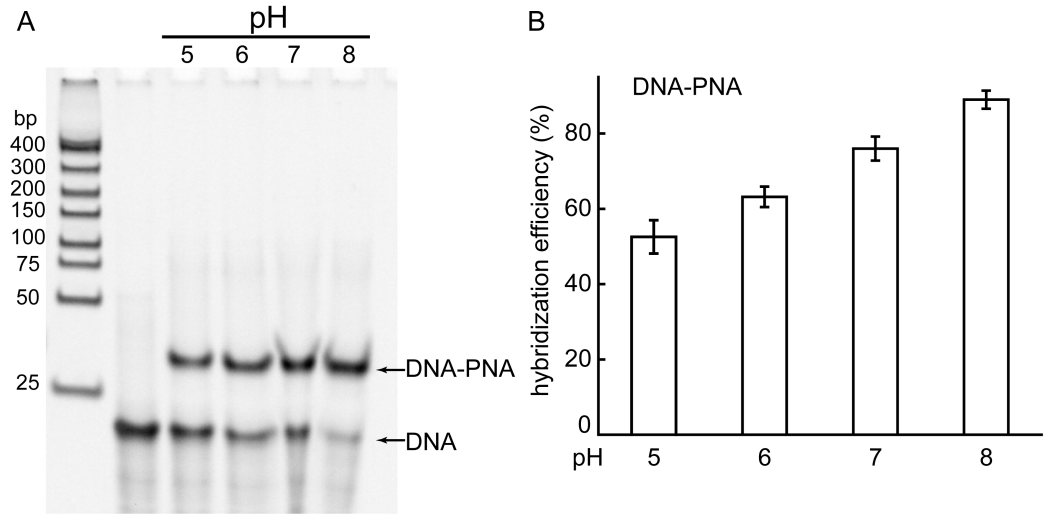

**Figure S7**. The hybridization efficiency between PNA/DNA with different pH. (A) Native 12% PAGE electrophoresis characterized the hybridization. (B) Gel quantitative analysis of the hybridization efficiency based on the band intensity of the different pH using the software BandScan.

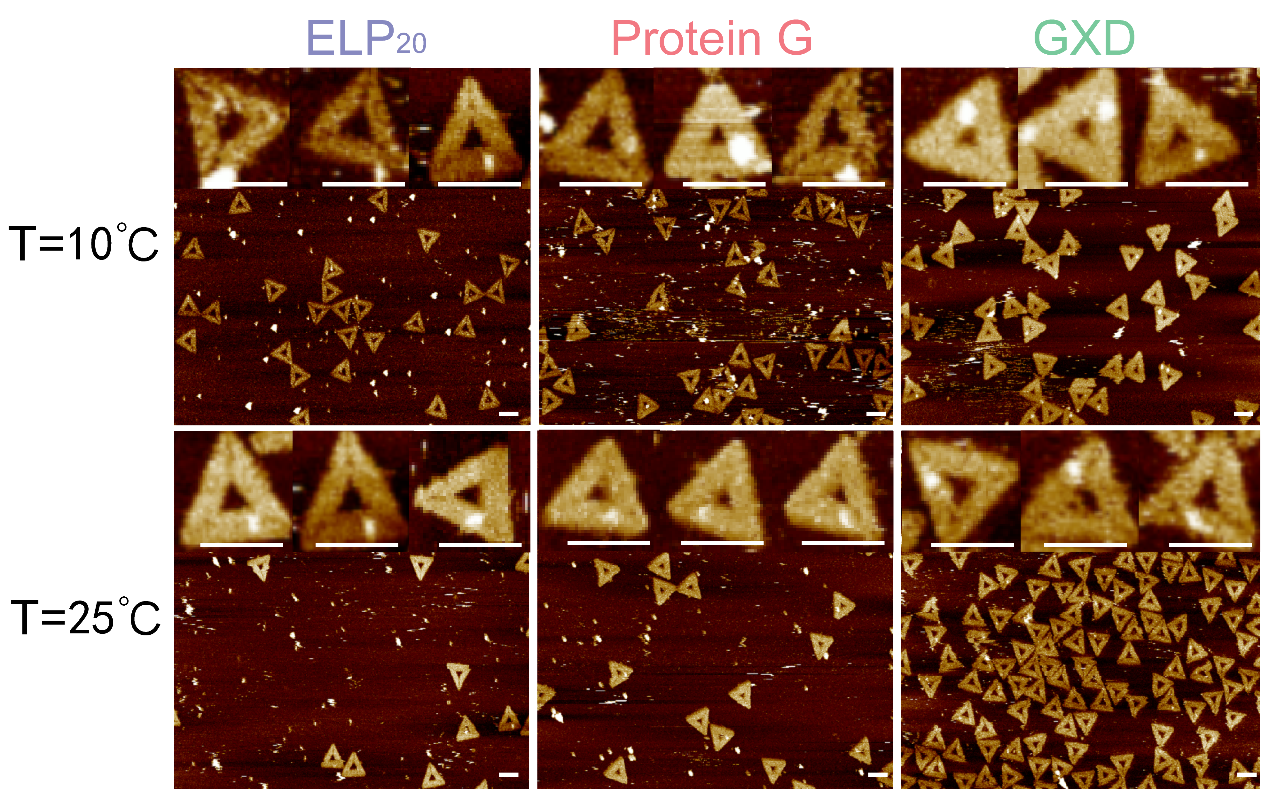

**Figure S8**. AFM images of site-specific organization of ELP_20_, protein G, and GXD through *Oa*AEP1 catalyzed ligation under different temperatures. All scale bars: 100 nm.

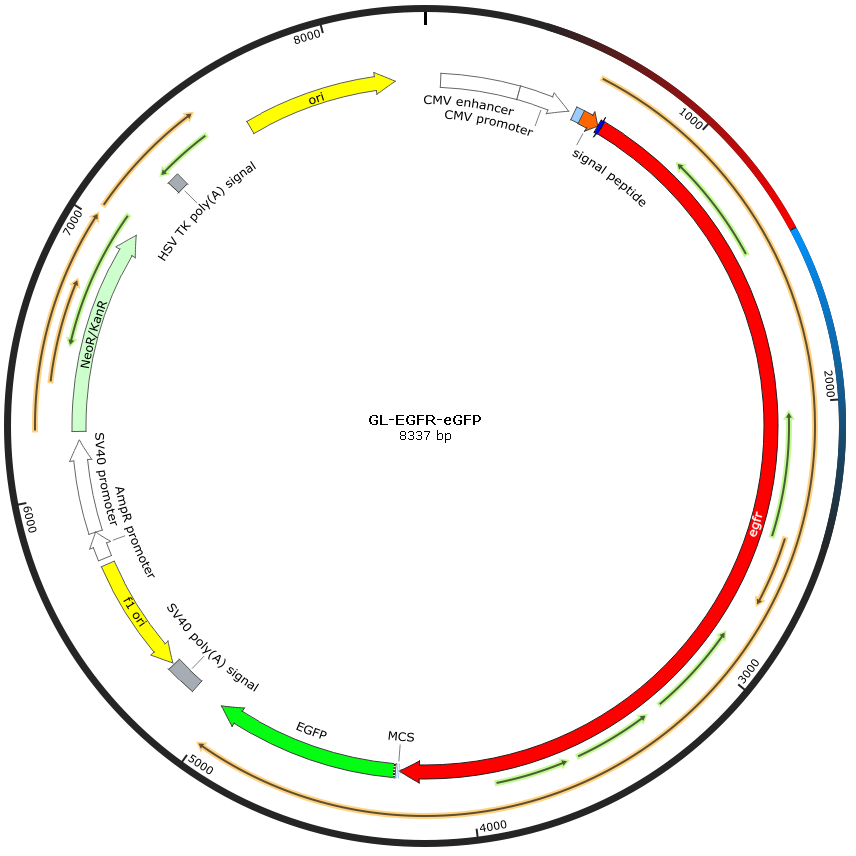

**Figure S9.** Vetor map of GL-EGFR-EGFP assembled by Gibson assembly (The first 24 amino acids of EGFR code for a signal peptide,which is cleaved off during biosynthesis. The DNA sequence of the GL-KGG peptide was therefore insertd after amino acid 24; between the Cterminal end of the EGFR signal peptide and N terminal end of the mature protein. After biosynthesis and cleavage of the signal peptide, the mature protein with N teminal GL-KGG-tag should be revealed). Image was made by Snap Gene Viewer 3.0.

Table S1. PNA sequences used in this study. Each PNA strand contains a linker (2- or 6-residue) followed by a NGL tripeptide at the C terminus.

| Name | Sequence (N→C) |
| --- | --- |
| PNA-2aa | ACTAGTACGCTC-**GlyGlyAsnGlyLeu** |
| PNA-6aa | ACTAGTACGCTC-**GlyGlySerGlyGlySerAsnGlyLeu** |

Table S2. Amino acid sequence of each POI. Each POI contains a GL dipeptide at the N terminus.

| Name | Sequence (N→C) |
| --- | --- |
| ELP | MGLHHHHHHGSVPGEGVPGVGVPGVGVPGVGVPGVGVPGAGVPGAGVPGGGVPGGGVPGEGVPGEGVPGVGVPGVGVPGVGVPGVGVPGAGVPGAGVPGGGVPGGGVPGEGRSC |
| Protein G | MGLHHHHHHPVSTTYKLVINGKTLKGETTTEAVDAATAEKVFKQYANDNGVDGEWTYDDATKTFTVTE |
| GB1-X module-Doc (GXD) | MGLHHHHHHGSMDTYKLILNGKTLKGETTTEAVDAATAEKVFKQYANDNGVDGEWTYDDATKTFTVTERSGGNTVTSAVKTQYVEIESVDGFYFNTEDKFDTAQIKKAVLHTVYNEGYTGDDGVAVVLREYESEPVDITAELTFGDATPANTYKAVENKFDYEIPVYYNNATLKDAEGNDATVTVYIGLKGDTDLNNIVDGRDATATLTYYAATSTDGKDATTVALSPSTLVGGNPESVYDDFSAFLSDVKVDAGKELTRFAKKAERLIDGRDASSILTFYTKSSVDQYKDMAANEPNKLWDIVTGDARS |

Table S3. Quantitative analysis of the reaction yields of POI-PNA conjugation under various conditions

|  | Yield (%) | | | | | | |
| --- | --- | --- | --- | --- | --- | --- | --- |
|  | Spacer length (aa)  (2 h, pH 7, 25 ℃) | | pH  (PNA-6aa, 2 h, 25 ℃) | | | Temp (℃)  (PNA-6aa, 2 h, pH 7) | |
|  | 2 | 6 | 5 | 6 | 7 | 10 | 25 |
| ELP_20_ | 34.2±2.9 | 41.2±4.3 | 22.7±2.5 | 26.5±3.9 | 47.7±5.4 | 38.9±3.2 | 48.4±2.8 |
| Protein G | 31.9±5.1 | 34.2±4.8 | 14.5±4.2 | 31.9±6.1 | 28.8±2.2 | 28.6±3.6 | 32.2±3.3 |
| GXD | 28.9±1.9 | 42.8±2.6 | 15.6±2.7 | 30.1±2.1 | 30.5±3.6 | 41.8±4.8 | 48.9±5.3 |

Table S4. Sequences used as capture strands in DNA origami nanostructure

| Name | Sequence (5’→3’) |
| --- | --- |
| Origami-capture-1 | GAGCGTACTAGTCTTTTTACCAGTCAGGACGTTGGAACGGTGTACAGACCGAAACAAA |
| Origami-capture-2 | GAGCGTACTAGTCTTTTTCCAAGCGCAGGCGCATAGGCTGGCAGAACTGGCTCATTAT |
| Origami-capture-3 | GAGCGTACTAGTCTTTTTACCTTATGCGATTTTATGACCTTCATCAAGAGCATCTTTG |

Table S5. DNA sequences used as probe strand and fuel strand in reversible protein labeling

| Name | Sequence (5’→3’) |
| --- | --- |
| DNA-32mer | GAGCGTACTAGTTTCACCGGCTTGAAGTGCCG |
| Cy5-DNA-26mer | Cy5-CGGCACTTCAAGCCGGTGTAATGGTC |
| Fuel strand-26mer | GACCATTACACCGGCTTGAAGTGCCG |
